# An end-to-end workflow for multiplexed image processing and analysis

**DOI:** 10.1101/2021.11.12.468357

**Authors:** Jonas Windhager, Bernd Bodenmiller, Nils Eling

**Affiliations:** Department of Quantitative Biomedicine, University of Zurich, Zurich, Switzerland; Institute for Molecular Health Sciences, ETH Zurich, Zurich, Switzerland; Life Science Zurich Graduate School, ETH Zurich and University of Zurich, Zurich, Switzerland

## Abstract

Simultaneous profiling of the spatial distributions of multiple biological molecules at single-cell resolution has recently been enabled by the development of highly multiplexed imaging technologies. Extracting and analyzing biologically relevant information contained in complex imaging data requires the use of a diverse set of computational tools and algorithms. Here, we report the development of a user-friendly, customizable, and interoperable workflow for processing and analyzing data generated by highly multiplexed imaging technologies. The *steinbock* framework supports image pre-processing, segmentation, feature extraction, and standardized data export. Each step is performed in a reproducible fashion. The *imcRtools* R/Bioconductor package forms the bridge between image processing and single-cell analysis by directly importing data generated by *steinbock*. The package further supports spatial data analysis and integrates with tools developed within the Bioconductor project. Together, the tools described in this workflow facilitate analyses of multiplexed imaging raw data at the single-cell and spatial level.

## 1 Introduction

Highly multiplexed imaging enables the simultaneous detection of tens of biological molecules (e.g., proteins, RNA), also referred to as “markers”, in their spatial tissue context. Recently established multiplexed imaging technologies rely on cyclic staining with immunofluorescently tagged antibodies [1, 2], or multiplexed staining with oligonucleotide-tagged [3, 4] or metal-tagged antibodies [5, 6]. The acquired data are commonly stored as multi-channel images, where each pixel encodes the abundances of all acquired markers at a specific position in the tissue. After data acquisition, bioimage processing and segmentation are conducted to extract data for downstream analysis. End-to-end multiplexed image analysis currently requires a diverse set of computational tools and complex analysis scripts.

Quantitative analysis of biological entities captured by multiplexed imaging requires processing of multi-channel images. This involves image extraction and pre-processing, image segmentation, and the quantification of biological objects such as cells. Multiplexed image segmentation is often performed by first classifying image pixels as nuclear, cytoplasmic or background (e.g., using software such as *Ilastik* [7]), followed by identifying and segmenting cells based on the resulting pixel-level class probabilities (e.g., using software such as *CellProfiler* [8]). Several pipelines have been developed for multi-channel image processing using pixel classification-based segmentation approaches, including the *IMC Segmentation Pipeline* [9], *imcyto* [10], and *MCMICRO* [11].

Classification-based segmentation approaches require the training of pixel classifiers (i.e., manual annotation of images), a process that is specific to the acquired markers. To enable applicability across marker panels, the dimensionality of input images can be reduced by aggregating selected channels. For example, a two-channel nuclear/cytoplasm image can be constructed from a multi-channel image by averaging all nuclear and all cytoplasmic channels. Such channel-aggregated images can then be used to train panel-agnostic pixel classifiers to imitate classifiers previously trained on specific sets of markers [12], or to directly apply generalist cell segmentation algorithms [13–15]. The latter methodologies include deep learning-enabled algorithms achieving human-level performance across various tissue types and imaging platforms [15].

In recent years, graphical user interface (GUI) software specialized for multiplexed imaging platforms have been developed to analyze cells with regards to their spatial location [16–19]. These tools are user-friendly and allow joint visualization of different data representations (e.g., images, single-cell features), but they often have little interoperability and are difficult to extend. Alternatively, after multi-channel image processing, the extracted tabular data can be analyzed using common programming languages such as R and Python. The *squidpy* Python package was developed recently to analyze spatial molecular data [20], and *giotto* performs similar analyses in R [21]. In contrast to stand-alone tools, the Bioconductor project offers interoperability among diverse analysis packages by relying on standardized data classes [22]. An example of such is the *SingleCellExperiment* class that supports general single-cell analyses including clustering of cells and dimensionality reduction [23–25], spatial clustering [26], and visualization of multiplexed imaging data [27].

## 2 Results

Here, we present a modular and interoperable computational workflow to process and analyze multiplexed imaging data. The *steinbock* framework facilitates multi-channel image processing including raw data pre-processing, image segmentation, and feature extraction. Data generated by *steinbock* can be directly read using the *imcRtools* R/Bioconductor package, which also provides functionality for data visualization and spatial analysis (Figure 1).

**Figure 1:**
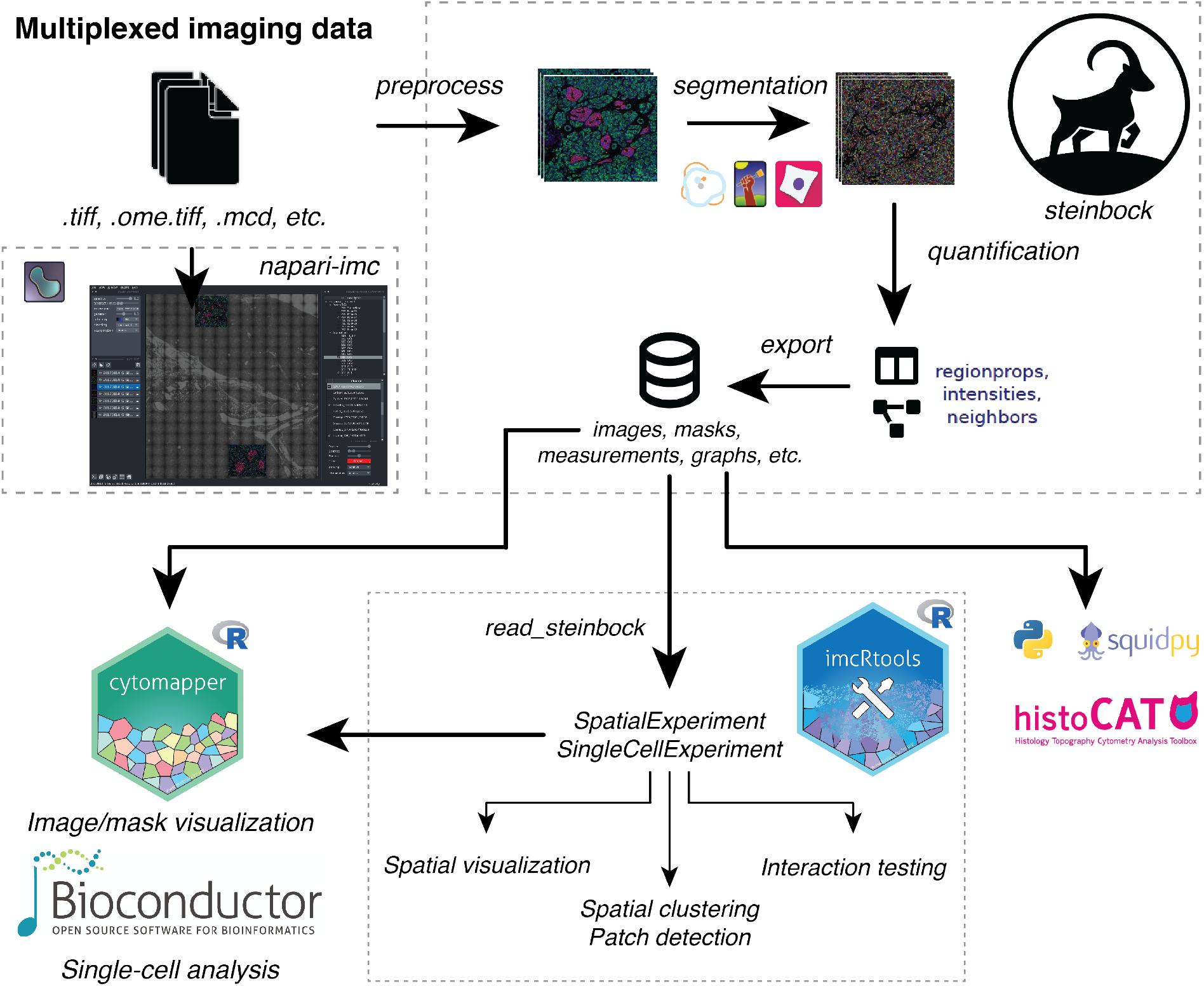
Overview of the multiplexed image processing and analysis workflow. Raw image data can be interactively visualized using *napari* plugins such as *napari-imc* for IMC to assess data quality and for exploratory visualization. The *steinbock* framework performs image pre-processing, cell segmentation and single-cell data extraction using established approaches and standardized file formats. Data can be read into R using the *imcRtools* package, which supports spatial visualization and analysis. Storing the data in *SingleCellExperiment* or *SpatialExperiment* objects, *imcRtools* integrates with a variety of data analysis tools of the Bioconductor project such as cytomapper [27]. Data can also be exported from *steinbock* as *anndata* objects to facilitate analysis in Python, e.g., using *squidpy* [20].

The presented workflow is user-friendly, customizable and reproducible, and integrates with a variety of downstream analysis strategies by employing standardized data formats. The tools included in this workflow support processing and analysis of data generated by a range of mul-tiplexed imaging technologies. For demonstration purposes, we present data from imaging mass cytometry (IMC), which relies on tissue staining with metal-labelled antibodies to simultaneously measure the spatial distribution of up to 40 proteins or RNA molecules at 1 µm pixel size [5, 28].

### 2.1 Multi-channel image visualization with napari

Visual inspection of imaging data is key to bioimage analysis, and specialized software is required for multi-channel image visualization [29, 30]. The recently developed multi-dimensional image viewer *napari* enables the fast and interactive visualization of multi-channel images, supported by a growing community of developers [31]. Through plugins written in Python, *napari* can be extended to load image data from a variety of multiplexed imaging platforms.

Multiplexed imaging often generates complex raw data that can be challenging to visualize and process. An example for such data is the proprietary MCD file format for IMC: After image acquisition, a single MCD file can hold raw acquisition data for multiple regions of interest, optical images providing a slide level overview of the sample (“panoramas”), and detailed metadata about the experiment. To facilitate IMC data processing, we created *readimc*, an open-source Python package for extracting the multi-modal (IMC acquisitions, panoramas), multi-region, multi-channel information contained in raw IMC files.

Building on *readimc*, we developed *napari-imc*, a modular plugin for loading raw IMC data into *napari*. Upon opening MCD files, *napari-imc* displays a graphical user interface for loading panoramas, acquisitions (Figure 2A) and channels (Figure 2B). For each loaded panorama and for each combination of loaded acquisition and channel, *napari-imc* creates an *image layer* (Figure 2C). In *napari*, image layers represent single-channel grayscale or color images that can be overlaid in the main panel (Figure 2D). Importantly, all image layers are spatially aligned. Adjusting channel settings (Figure 2E) will broadcast the chosen values to the settings of all associated image layers (Figure 2F).

**Figure 2:**
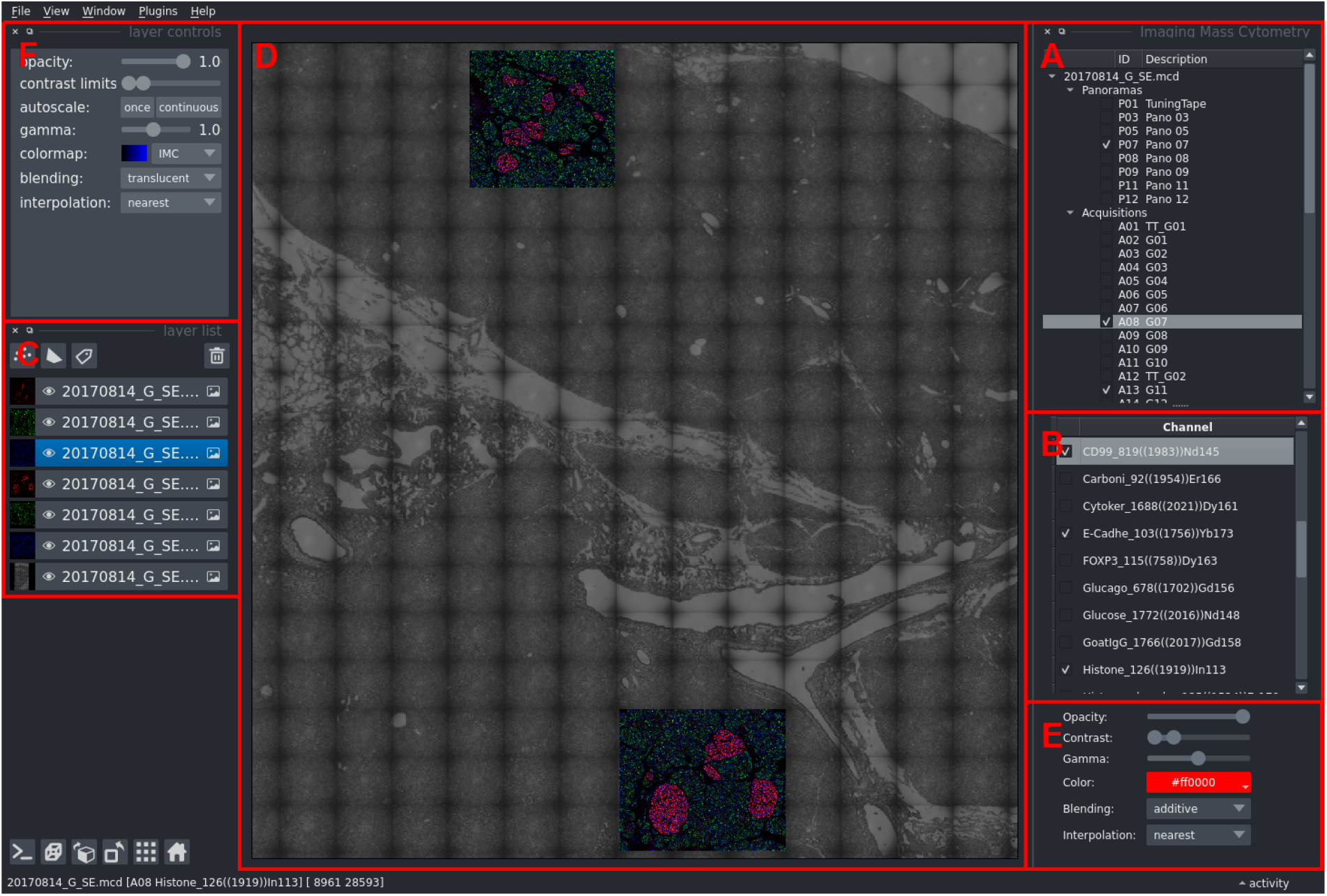
Visualization of raw IMC data using napari-imc. Screenshot of *napari-imc* visualizing two selected acquisitions from Damond et al. [32] overlaid on one panorama. Image channels corresponding to markers for CD99 (red, cytoplasm), E/P-Cadherin (green, cytoplasm), and Histone H3 (blue, nucleus) are displayed. Labeled boxes (red) indicate GUI elements of *napari* (C,D,F) and *napariimc* (A,B,E): **(A)** MCD files containing panoramas and multi-channel acquisitions, **(B)** image channels of all loaded acquisitions, **(C)** image layers corresponding to loaded panoramas or per-channel images from individual IMC acquisitions, **(D)** visualization of the image layers, **(E)** settings of the selected image channel that are broadcast to the properties of associated image layers, **(F)** properties of the image layers.

The *napari-imc* plugin enables the rapid visualization and quality control of raw IMC data, without the need for manual file conversion. Unlike existing software, *napari-imc* loads multi-channel acquisitions and panoramas into a shared coordinate system, allowing for quick spatial orientation with respect to the physical tissue slide. Jointly configuring layers by their channel enables the qualitative comparison of multiple regions of interest. Further, owing to its modular implementation, *napari-imc* is ready for extension to similar data formats in the future. In summary, using *napari-imc*, the user can perform a quick, qualitative inspection of multiplexed image acquisitions prior to downstream image processing.

### 2.2 Multi-channel image processing with steinbock

Image processing forms the basis for data analysis in any quantitative imaging project. Core tasks of image processing include extraction of images from raw data, segmentation and quantification of spatial entities such as cells, and data export for downstream analysis. Despite their repetitive character, each step requires meticulous quality control of its intermediate outputs, a characteristic often neglected by fully automated image processing pipelines. Further, the individual processing steps must be reproducible, and software must be easy to install and use. To ensure compatibility, all inputs and results must interface with existing tools and approaches.

To facilitate multiplexed image analysis, we developed *steinbock*, a collection of tools for rapid processing of multi-channel images. The *steinbock* framework builds on existing approaches, and can be operated through its easy-to-use *command-line interface (CLI)*. Using simple commands, multi-channel images can be processed step by step, offering full control of the individual tasks. The framework is distributed as a *Docker container* that bundles third-party software required for the individual tasks and offers platform-independence and reproducibility. Internally, *steinbock* is implemented in Python, and the *steinbock Python package* can be used programmatically.

Both the *steinbock* CLI and the *steinbock* Python package are fully documented and have been extensively tested. The *steinbock* framework integrates with existing software by processing data in standardized formats and its modular open-source implementation enables community-driven development. The tools provided by *steinbock* can be used to build multiplexed image processing workflows, a typical example is shown in Figure 3.

**Figure 3:**
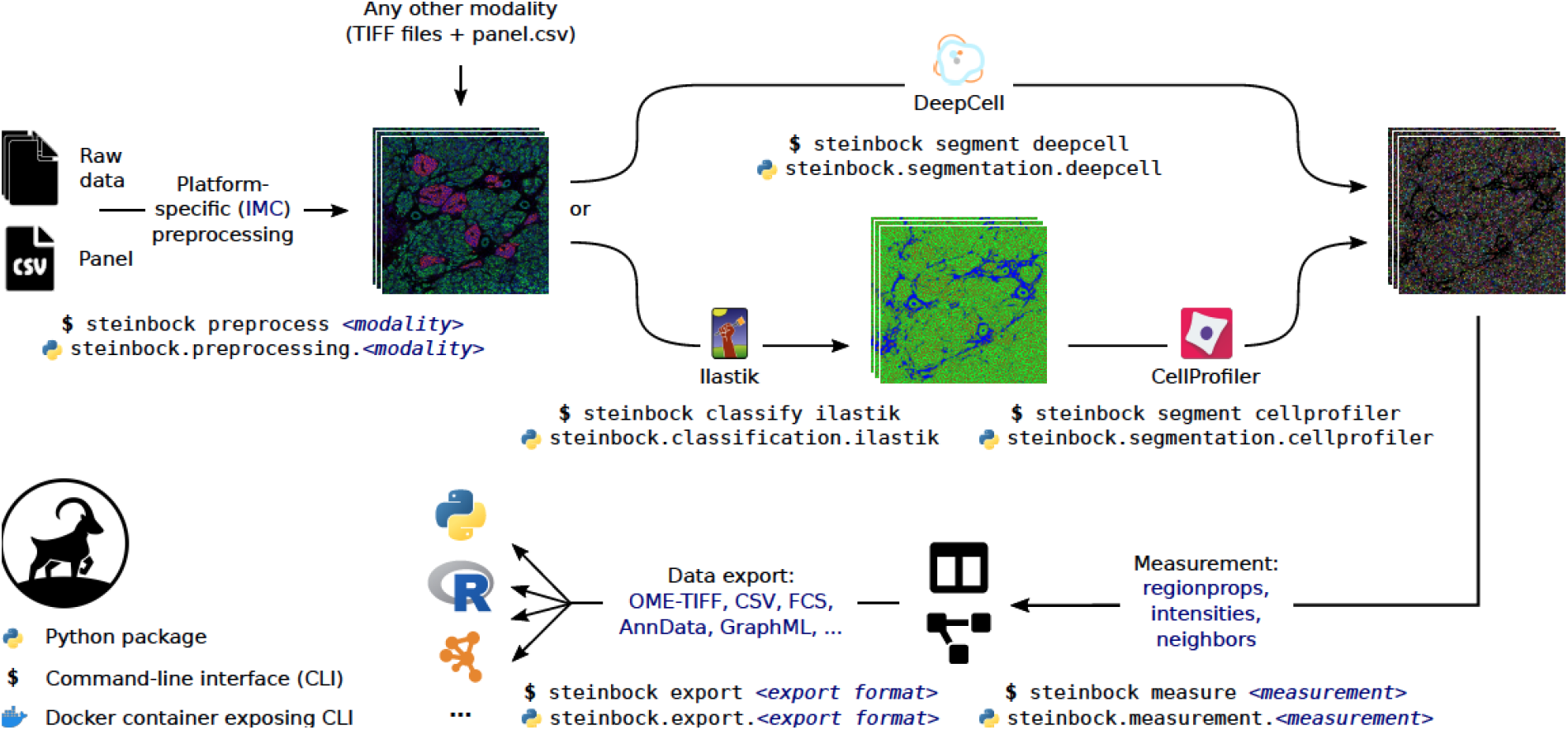
A typical multiplexed image processing workflow using steinbock. Individual steps are indicated as arrows, with corresponding *steinbock* commands and Python modules shown below. Further parameters and third-party software (bundled in the *steinbock* Docker container) are overlaid on the arrows. Input and output data are indicated conceptually and described in further detail in Section 2.2.

#### 2.2.1 Data input and pre-processing

The *steinbock* framework operates on multi-channel TIFF images, which can be either provided by the user or extracted from supported raw data formats using *steinbock*. When starting from supported raw data formats, images are extracted and pre-processed on a pixel level tailored to the specific imaging platform. In the case of IMC, this involves image extraction from raw data using *readimc*, extraction of metadata, filtering and sorting of image channels, and removal of hot pixels as described in the original *IMC Segmentation Pipeline* [9]. To enable processing of large images that would otherwise exhaust computational resources, *steinbock* offers functionality for tiling and stitching images.

In addition to the images, *steinbock* requires channel-specific metadata (e.g., channel names), which can be supplied in CSV format (e.g., a “panel file”). In the case of IMC, such panel files can be automatically generated from the raw data.

#### 2.2.2 Image segmentation

After image extraction, image segmentation is performed to define spatial entities such as cells, yielding segmentation masks (e.g., cell masks). Segmentation masks are single-channel images that match the input images in size, with non-zero grayscale values indicating the IDs of segmented objects. The *steinbock* framework explicitly supports the following supervised image segmentation approaches:

- **Random forest-based** image segmentation as presented by Zanotelli and Bodenmiller [9]. Briefly, a random forest is trained using *Ilastik* [7] on randomly extracted image crops and selected image channels to classify pixels as nuclear, cytoplasmic, or background. Employing a customizable *CellProfiler* [8] pipeline, the probabilities are then thresholded for segmenting nuclei, and nuclei are expanded into cytoplasmic regions to obtain cell masks.
- **Deep learning-based** image segmentation as presented by Greenwald et al. [15]. Briefly, *steinbock* first aggregates user-defined image channels to generate two-channel images representing nuclear and cytoplasmic signals. Next, the *DeepCell* Python package is used to run *Mesmer*, a deep learning-enabled segmentation algorithm pre-trained on *TissueNet* [15], to automatically obtain cell masks without any further user input.

#### 2.2.3 Object quantification

Following image segmentation, features of the detected objects (e.g., cells) are quantified. The *steinbock* framework is equipped with functionality for measuring region properties (e.g., area, eccentricity), aggregated marker intensities (e.g., mean, median), and spatial neighbors. The measurement of spatial neighbors yields *spatial object graphs*, in which nodes correspond to objects, and nodes in spatial proximity are connected by an edge. Distances between objects are computed based on the objects’ centroids or borders. These distances are used to construct spatial object graphs by distance thresholding or *k*-nearest neighbor (*k*-NN) detection. Additionally, *steinbock* can construct spatial object graphs by the means of morphological dilation (“pixel expansion”). The choice of neighborhood measurement depends on the downstream data analysis approach. For example, pixel expansion is commonly used for neighborhood analysis as presented by Schapiro et al. [16].

#### 2.2.4 Data output and export

All data generated by *steinbock* are stored in standardized, well-documented file formats that are directly supported by a wide range of third-party software. Specifically, the *imcRtools* package can be used to load single-cell data generated by *steinbock* into *SingleCellExperiment* [25] or *SpatialExperiment* [33] objects for downstream analysis in R (described in detail in Section 2.4).

To further facilitate compatibility with downstream analysis, data can additionally be exported to a variety of file formats such as OME-TIFF for images, CSV and FCS for single-cell data, the *anndata* [34] format for data analysis in Python, and various graph file formats for network analysis using software such as *CytoScape* [35]. For export to OME-TIFF, *steinbock* uses *xtiff*, a Python package we developed for writing multi-channel TIFF stacks.

### 2.3 Segmentation quality control with cytomapper

Visual assessment of image segmentation quality is crucial to avoiding biases in downstream analyses. Visualization is commonly done by outlining segmented objects (e.g., cells) on composite images showing features of interest (e.g., marker proteins). We previously developed the *cytomapper* R/Bioconductor package [27] to visualize multi-channel images and to map cellular features onto segmentation masks. The *cytomapper* package supports reading and storage of multi-channel images and segmentation masks including the TIFF files generated by *steinbock* (Figure 1). Upon data import, multi-channel images can be visualized as composite images with up to six colors (Figure 4A). More importantly in the context of image segmentation, multi-channel images and segmentation masks can be combined to outline segmented cells on composite images (Figure 4B).

**Figure 4:**
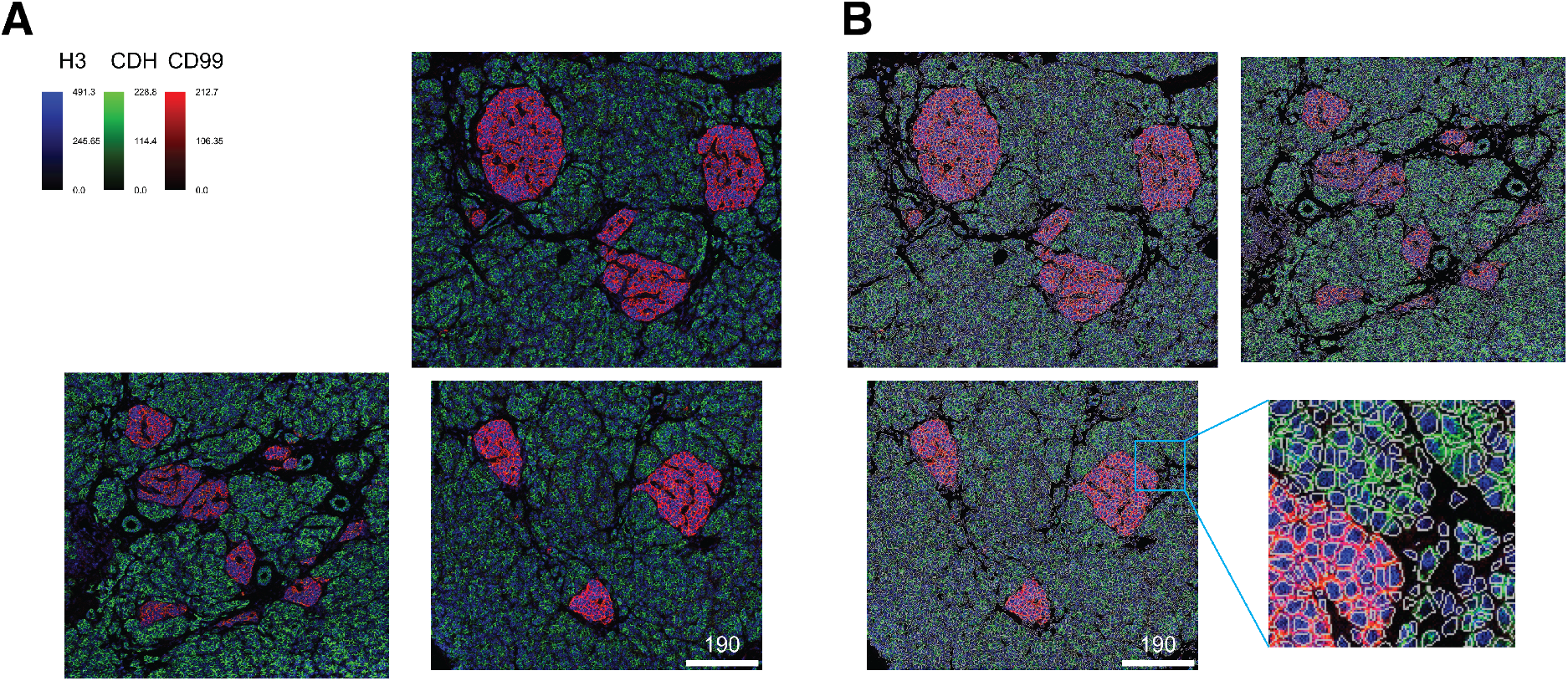
Visualization of multiplexed images and segmentation masks using cytomapper. **(A)** Composite images showing Histone H3 (H3, blue, nucleus), E/P-cadherin (CDH, green, cytoplasm), and CD99 (red, cytoplasm) expression. **(B)** Composite images from (A) with segmented cells outlined in white with a zoom plot shown to the right.

### 2.4 Spatial data analysis with imcRtools

Highly multiplexed imaging generates images from which, after image segmentation, object-specific measurements can be extracted. Most often, segmented objects represent cells in their tissue context. Common tasks for single-cell data analysis include clustering and cell phenotyping, dimensionality reduction, and differential analysis; standardized methods for these tasks have been previously developed [25]. The *imcRtools* R package, available through the Bioconductor project, provides functionality that supports handling of data derived from multiplexed imaging technologies, spatial single-cell analysis, and spatial visualization.

During image pre-processing, *steinbock* generates text files containing the summarized intensity per cell and channel, properties of the segmented cells (e.g., location, morphology), and spatial object graphs indicating cells in close proximity. The *imcRtools* package reads the *steinbock* output and jointly stores single-cell and neighborhood information in a *SpatialExperiment* or *SingleCellExperiment* object (Figure 1). These classes directly support single-cell analysis using a variety of analysis packages, including *scater* [24], *scran* [23], *BayesSpace* [26], and *cytomapper* [27]. The following sections describe functions provided by *imcRtools* that take a *SpatialExperiment* or *SingleCellExperiment* object as standardized input.

#### 2.4.1 Spatial data visualization

The visualization of multi-channel images and mapping of cellular features onto segmentation masks can be performed using the *cytomapper* package [27]. Complementary to this pixel-level visualization strategy, the *imcRtools* package supports visualizing locations of cells as well as *spatial object graphs*. The centroids of cells are visualized as points, and lines are drawn between cells in spatial proximity (also referred to as neighboring cells). The lines represent edges in the *spatial object graphs* computed by *steinbock* or *imcRtools*. The *imcRtools* package allows coloring of cells based on marker expression or cellular metadata (e.g., cell type, morphological features; Figure 5A). The size and shape of points can also be adjusted based on cellular metadata. Lines between neighboring cells can be colored by edge features or cellular metadata associated with the cell from which the edge originates (Figure 5B).

**Figure 5:**
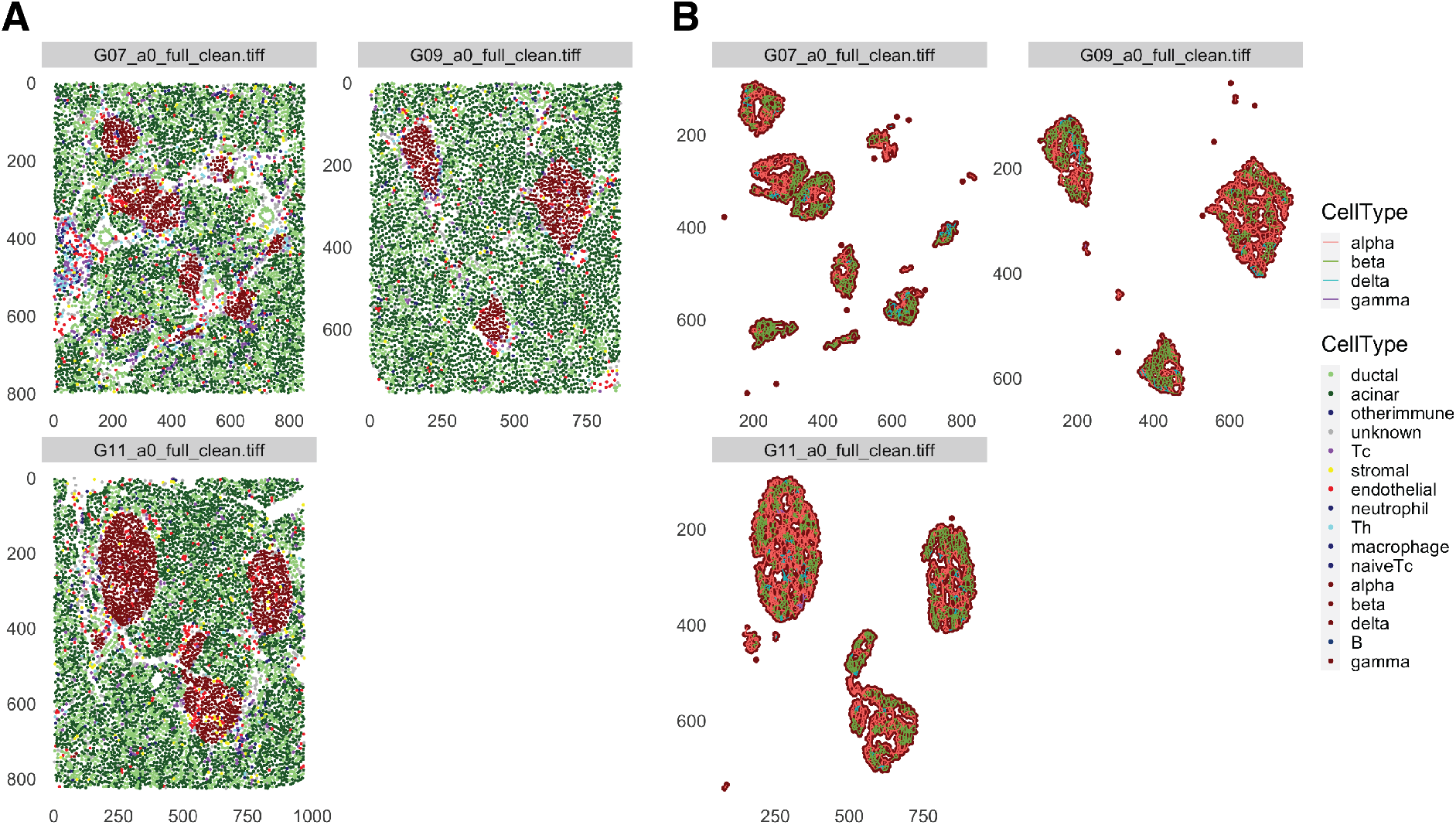
Spatial visualization of cells and their neighborhood information. **(A)** Points represent the centroids of segmented cells and are colored based on the cells’ phenotypes as defined in the original publication [32]. Cells from images displayed in Figure 4 were selected. Islet cells are shown in red; immune cells except T helper (Th) cells and cytotoxic T (Tc) cells are shown in dark blue. **(B)** A *k*-NN spatial object graph was constructed with *k* = 5. Only pancreatic islet cells are visualized. As in (A), points represent the centroids of segmented cells and are colored by their phenotype. Cells in spatial proximity are visualized as lines (i.e., edges) between points. Lines are colored by the phenotypes of the cells from which edges originate.

#### 2.4.2 Spatial data analysis

Single-cell resolved multiplexed imaging produces data that allows analysis of cells with regards to their spatial location. Over the past five years, spatial data analysis approaches have been developed that yield biologically meaningful insights from highly multiplexed imaging data [4, 16, 28, 36, 37]. The *imcRtools* package supports such data analysis approaches in a standardized fashion within the Bioconductor framework.

The *imcRtools* package can be used to summarize each cell’s neighborhood by aggregating either across cellular metadata or the expression of neighboring cells. Cells can now be clustered based on aggregated values of neighboring cells, an approach that has been previously proposed by Goltsev et al. [4] and Schürch et al. [36] to characterize cellular neighborhoods. To demonstrate the neighborhood aggregation approach with subsequent clustering, we used *imcRtools* to analyze a type 1 diabetes dataset provided by Damond et al. [32] as part of the *imcdatasets* R/Bioconductor package [38]. For each cell, the proportion of cell types among the 10 nearest neighbors were computed. This information was subsequently used to cluster all cells. The spatial clusters detected in this manner represent different tissue compartments: spatial clusters 1, 4, and 5 contain islet cells, and spatial clusters 3, 6, 7, 8, and 9 include mainly cells from exocrine tissue. Within the latter, clusters 6, 7, and 9 contain ductal and acinar cells, and clusters 3 and 9 contain a mix of exocrine and stromal cells. Spatial cluster 10 is mainly composed of T cells and endothelial cells, and cluster 2 involves aggregates of B and T cells (Figure 6A). Interestingly, spatial cluster 2 appears in proximity to islets in images obtained from early onset diabetes patients (Figure 6B), possibly indicating immune cell and islet cell interactions.

**Figure 6:**
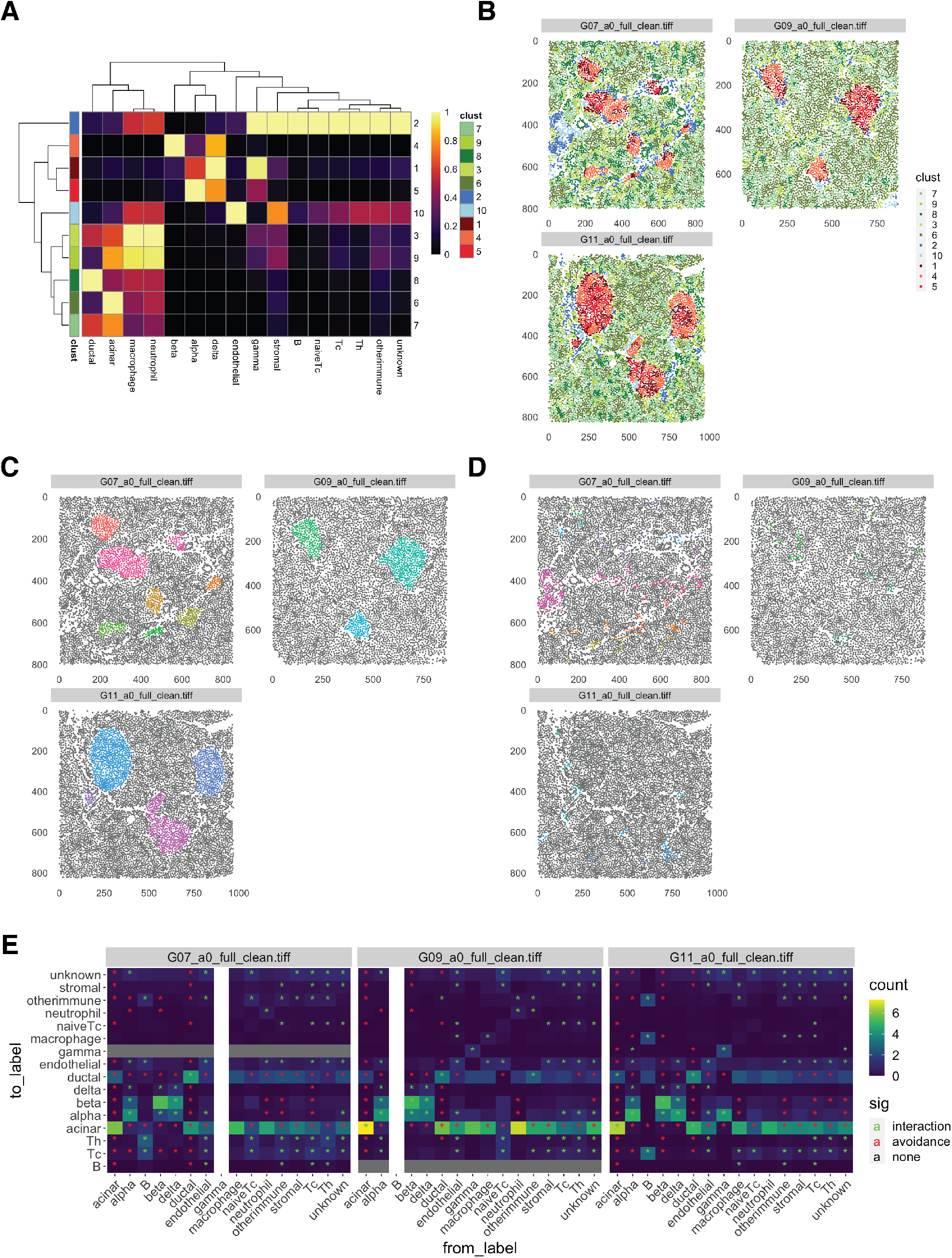
Spatial analysis strategies supported by imcRtools. **(A)** Cells of the type 1 diabetes dataset generated by Damond et al. [32] were clustered based on cell type proportions among their 10 nearest neighbors. Cell type proportions within each spatial cluster (rows, colored by cluster) are shown as a heatmap. Columns are rescaled between 0 and 1. **(B)** Points represent the centroids of segmented cells and their color indicates spatial clusters. Cells from images displayed in Figure 4 were selected. **(C)** *α, β, δ*, and *γ* cells were selected to detect connected patches of islet cells and patches were expanded by 10 µm. Colored points represent cells within islet patches and grey points indicate cells outside islet patches. **(D)** Immune cells were selected to detect immune cell aggregates. Colored points represent cells within immune patches and grey points indicate cells outside immune patches. **(E)** The interaction count between pairs of cell types visualized in form of one heatmap per image. Cell types that interact more often than expected by chance are indicated by a green star (interaction). Cell types that interact less often are indicated by a red star (avoidance). Statistical significance was defined at an empirical p-value threshold of 0.01.

As an alternative approach, the algorithm developed by Hoch et al. can be used to detect patches [37], where patches are defined as cells of a specific phenotype in spatial proximity. In an optional step, a concave or convex hull is built around the detected patches. This hull can be expanded to include cells in direct vicinity as demonstrated for islet cell types (Figure 6C) and immune cell types (Figure 6D).

The *imcRtools* package further allows statistical testing of whether cell types interact more or less frequently than expected by chance using a method implemented by Schapiro et al. as part of the *histoCAT* software [16]. In this context, interacting cells are defined as cells in spatial proximity (i.e., neighboring cells detected by constructing a spatial object graph). First, for each cell of type A the number of neighbors of type B are counted. In each image, this count is averaged in one of three different ways, depending on the question at hand:

1. The count is divided by the total number of cells of type A. The final count can be interpreted as “How many neighbors of type B does a cell of type A have on average?”.
2. The count is divided by the number of cells of type A that have at least one neighbor of type B. The final count can be interpreted as “How many neighbors of type B has a cell of type A on average, given it has at least one neighbor of type B?”
3. For each cell, the count is binarized to 0 (less than a specified number of neighbors of type B) or 1 (more than or equal to a specified number of neighbors of type B). The binarized counts are averaged across all cells of type A. The final count can be interpreted as “What fraction of cells of type A have at least a given number of neighbors of type B?”. This counting strategy was proposed by Schulz et al. [28].

Next, the computed count is compared to an empirical null distribution of interaction counts. To derive such a distribution, cell labels are randomized a number of times, and for each iteration the interaction count is computed. Statistical inference is performed by comparing the actual interaction count to the empirical null distribution.

As an example, we analyzed the type 1 diabetes data reported by Damond et al. [32]. For all three example images, we observed that *α, β*, and *δ* cells interact more often than expected when cell types are randomly distributed across the images (Figure 6E). This result was expected, because *α, β*, and *δ* cells are locally confined in pancreatic islets. In addition, we observed increased interactions between naïve cytotoxic T cells (naïve Tc), cytotoxic T cells (Tc) and helper T cells (Th) (Figure 6E).

As these examples demonstrate, the *imcRtools* package provides functionality to (i) handle data extracted from multiplexed images, (ii) visualize cellular information in a spatial manner and (iii) perform spatial data analysis to detect cell aggregates and enriched interactions between cell types. Building upon standardized data classes, the *imcRtools* package connects bioimage processing as performed using *steinbock* with downstream analyses supported by the Bioconductor project.

## 3 Discussion

Here, we present a modular workflow for analyzing highly multiplexed imaging data, introducing the *steinbock* framework for multi-channel data processing and *imcRtools* for spatial data analysis. The workflow standardizes common processing and data analysis tasks and integrates bioimage and single-cell analysis in a user-friendly fashion. Due to its modular structure, the workflow is easily extendable to additional image processing steps and new approaches for single-cell and spatial analysis.

Starting from multi-channel TIFF files or raw data from supported imaging platforms, the *steinbock* framework extracts data for downstream analysis. Depending on the imaging platform, additional pre-processing, such as the alignment of consecutive images for cyclic immunofluorescence modalities [1, 2], may be required before *steinbock* is run. For IMC, however, *steinbock* can be directly applied to raw acquisition data, and it performs established pre-processing routines [9]. Explicit support for other imaging platforms may be added in the future. Furthermore, processing of large images is enabled by image tiling/stitching. As with all bioimaging approaches, meticulous quality control should be performed at each step of the image processing workflow, including careful inspection of the raw imaging data using tools such as *napari* [31].

A core task of multi-channel image processing is the segmentation of spatial entities such as cells. The *steinbock* framework currently supports two previously published image segmentation approaches [9, 15]. The first is the random forest-based approach [9], which is highly customizable, allows for dataset-specific tuning, and has been applied in several multiplexed imaging applications [28, 32, 39]. By adapting the provided *Ilastik* [7] and *CellProfiler* [8] pipelines, this approach enables segmentation of arbitrary spatial entities such as tumor/stroma regions or pancreatic islets. However, the random forest classifier needs to be manually trained for each dataset and is limited in its applicability to other projects. The second approach is based on deep learning [15], which builds on existing annotations [15] and can be used to rapidly segment cells or nuclei without the need for manually training a classifier. Irrespective of the chosen approach, the validity of the obtained segmentation masks always needs to be verified using tools such as *cytomapper* [27]. Further approaches for multi-channel image processing may be added in the future.

Unlike most existing software, *steinbock* was not implemented as a pipeline, but as a collection of independent, easy-to-use tools. Although these tools can be employed in streamlined pipelines, they are intended to be used in step-by-step workflows that offer full control of the individual tasks. This is reflected in the design principles underlying the *steinbock* framework, which focus on usability and interoperability with external tools. By operating on standardized data formats, *steinbock* integrates with the growing landscape of multiplexed imaging software and facilitates exploratory bioimage analysis.

To bridge bioimage processing, segmentation and feature extraction with downstream analysis strategies commonly performed in R, we developed the *imcRtools* package. The *imcRtools* package offers analysis strategies applicable to single-cell data extracted from most multiplexed imaging technologies [1–6] and can read in data generated by *steinbock*. It also supports spatial analysis and visualization of single-cell data derived from spatial transcriptomic technologies such as *MERFISH* [40] and *seqFISH* [41]. Furthermore, the *imcRtools* package contains functionality specifically for handling and processing of IMC data. These include reading in raw IMC data into *CytoImageList* objects for visualization with *cytomapper* or into *SingleCellExperiment* objects to perform channel spillover correction using the *CATALYST* R/Bioconductor package [42].

As part of the Bioconductor project [22] building upon the *SingleCellExperiment* [25] and *SpatialExperiment* [33] data classes, the *imcRtools* package fully integrates with a variety of single-cell and spatial analysis approaches and tools. It furthermore standardizes analysis approaches that were previously developed for highly multiplexed imaging data [4, 16, 28, 36, 37] and therefore complements other spatial data analysis tools such as *giotto* [21] and spatial clustering approaches including *BayesSpace* [26] and *lisaClust* [43]. As highly multiplexed imaging technologies and new ways of interpreting spatially annotated data develop, the *imcRtools* package will provide a platform for implementing emerging data analysis strategies.

## 4 Methods

### 4.1 Example data

To highlight the functionality of the *imcRtools* and *cytomapper* R/Bioconductor packages, an IMC dataset containing pancreatic islets of healthy and type 1 diabetes patients was reanalyzed [32]. The data are available in the *imcdatasets* R/Bioconductor package [38] (version 1.2.0) under the accessor: *DamondPancreas2019Data*.

### 4.2 Multi-channel image visualization

Raw IMC data from Damond et al. [32] was visualized using *napari* 0.4.11 and *napari-imc* 0.6.2 for illustration purposes (Figures 2, 3). Two acquisitions in spatial proximity were selected and overlaid on the corresponding panorama (Figure 2), matching the acquisitions in downstream analyses. Channels were configured to match Figure 4, applying the same channel-specific settings to all image layers.

### 4.3 Multi-channel image processing

The *steinbock* framework was implemented in the *steinbock* Python package (version 0.10.0), with Python modules corresponding to individual image processing tasks and including a commandline interface (CLI). The *steinbock* Docker container bundles the *steinbock* Python package with third-party software required for the individual tasks and exposes the *steinbock* CLI as Docker entrypoint.

The pre-processing module of *steinbock* currently supports reading raw IMC data using *readimc* 0.4.2 (developed in-house). Random forest-based image segmentation was implemented using *Ilastik* 1.3.3post3 [7] and *CellProfiler* 4.2.1 [8], and HDF5 files are used for loading multi-channel images in *Ilastik*. Deep learning-based image segmentation was implemented using *DeepCell* 0.11.0 [15], currently supporting *Mesmer* [15] for nuclear and cell segmentation. Further details on object quantification, data output, and data export can be found in the online documentation.

### 4.4 Segmentation quality control

#### Image visualization with cytomapper

##### All R based data analyses was performed using R version 4.1.2

The *cytomapper* package version 1.6.0 was used to generate composite images and to overlay segmentation masks on composite images using the *plotPixels* function to generate Figure 4.

The three largest images considering the number of pixels were selected from the dataset. The channel contrast was increased by scaling the intensities with a factor of 10 for Histone H3, 5 for E-/P-cadherin and 5 for CD99.

### 4.5 Spatial data analysis

#### Reading in the steinbock data

The *imcRtools* package provides the *read steinbock* function that reads in the summarized intensities per cell and channel, properties of the segmented cells (e.g., location, morphology), and spatial object graphs indicating cells in close proximity. These information are stored in either a *SpatialExperiment* [33] or a *SingleCellExperiment* [25] object. The cell intensity data are stored in the *counts* assay slot, object properties are stored in the *colData* slot, and the spatial interaction graphs are stored in a *colPair* slot. In the case of the *SpatialExperiment* object, spatial coordinates are stored in the *spatialCoords* slot, and spatial coordinates in a *SingleCellExperiment* are stored in the *colData* slot.

The *imcRtools* package contains functionality applicable across different multiplexed imaging technologies. In addition, *imcRtools* further supports IMC-specific data handling. Raw IMC data can be read into *CytoImageList* objects for visualization with *cytomapper*. Additionally, raw .txt files from control acquisitions can be read into a *SingleCellExperiment* object for spillover estimation using *CATALYST* [42].

#### Spatial graph construction and visualization

The graph construction, spatial visualization, and spatial analysis presented here were performed using *imcRtools* version 1.0.0. To visualize the location of cells and their interactions in Figure 5, a *k*-NN graph based on the cells’ centroids was constructed for *k* = 5 using the *buildSpatialGraph* function from the *imcRtools* package. The *buildSpatialGraph* function additionally supports constructing *spatial object graphs* based on distance thresholding or Delaunay triangulation to identify cells in close spatial proximity. The *plotSpatial* function was used to visualize the cells’ locations as points and to draw lines between cells if they were detected as neighbors using the 5-nearest neighbor graph construction approach.

#### Spatial data analysis

For the unsupervised spatial clustering analysis presented in Figure 6A,B, a *k*-NN graph based on the cells’ centroids was constructed for *k* = 10 using the *buildSpatialGraph* function. The *aggregateNeighbors* function from the *imcRtools* package was used to calculate the proportions of cell types among the 10-nearest neighbors. Cells were further clustered based on these proportions using k-means clustering with *k* = 10. For each spatial cluster, the number of cells of each cell type was divided by the cluster size. Next, fraction values were rescaled between 0 and 1 per cell type (Figure 6A). Instead of aggregating the cell type, the mean or median expression across all neighboring cells can be computed using the *aggregateNeighbors* function.

Figure 6C,D present an alternative approach to detect spatial aggregates (i.e., patches) of cells. The *imcRtools* package provides the *patchDetection* function, which detects patches of neighboring cells of a pre-defined type. To detect the pancreatic islets (Figure 6C), a 10-nearest neighbor graph was constructed using the cells’ centroids, and neighboring *α, β, δ* and *γ* cells were selected to detect connected patches. A concave hull was constructed around individual patches, and the hull was expanded by 10 µm to include cells in direct vicinity. Next, to detect patches that define interacting immune cells (Figure 6D), the 10-nearest neighbor graph was used to find patches of neighboring Tc, Th, naïve Tc, neutrophiles, macrophages, other immune cells and B cells. Patches with at least 4 cells were considered to represent aggregations of immune cells.

To test whether cell types interact more or less frequently compared to a random distribution, the *testInteractions* function available in the *imcRtools* package was used. First, a 10-nearest neighbor graph was constructed to detect neighboring cells. For each cell type pair “A” and “B”, the overall interaction count was divided by the number of cells of type “A” (Figure 6E). The count is compared to the random distribution of interaction counts, which was derived by permuting cell type labels 1000 times. Statistical significance for avoidance was defined if more than 990 iterations of random permutations produced larger counts. Statistical significance for association was defined if more than 990 iterations of random permutations produced smaller counts.

## 5 Software and code availability

The *readimc* Python package is available from https://github.com/BodenmillerGroup/readimc and installable via *pip*. Documentation is available at https://bodenmillergroup.github.io/readimc.

The *napari-imc* Python package/*napari* plugin is available from https://github.com/BodenmillerGroup/napari-imc and installable via *pip*, from within *napari* or from https://www.napari-hub.org.

The *steinbock* framework is available from https://github.com/BodenmillerGroup/steinbock. The *steinbock* Python package is installable via *pip*. The *steinbock* Docker container can be obtained via Docker from ghcr.io/bodenmillergroup/steinbock. Documentation is available at https://bodenmillergroup.github.io/steinbock

The *xtiff* Python package is available from https://github.com/BodenmillerGroup/xtiff and installable via *pip*.

The *imcRtools* development version is available from https://github.com/BodenmillerGroup/imcRtools. The release version is installable via Bioconductor: https://bioconductor.org/packages/imcRtools. Documentation is available at https://bodenmillergroup.github.io/imcRtools/.

The images and masks shown for illustration purposes in Figure 3 were taken from publicly available resources [32] and visualized using Python and *napari*. For convenience of the reader, the corresponding Jupyter notebook for loading multi-channel images, cell masks and cell outlines in *napari* is available from https://github.com/BodenmillerGroup/IMCDataAnalysis/tree/biorxiv_submission/code/mask_overlay.

The R analysis code used to highlight the functionality of the *cytomapper* and *imcRtools* packages is available from https://github.com/BodenmillerGroup/IMCDataAnalysis/tree/biorxiv_submission.

## 6 Contributions

J.W. developed and maintains the *steinbock* framework, *napari-imc, readimc* and *xtiff*. N.E. developed and maintains the *imcRtools* R package. J.W. and N.E. wrote the manuscript. B.B. carried senior authorship responsibility. All authors approve the manuscript.

## 7 Acknowledgments

We want to thank Jana R. Fischer, Daniel Schulz and Tobias Hoch for code contributions to the *imcRtools* package. Special thanks go to Vito R.T. Zanotelli for code contributions to the *imcRtools* package and for developing the conceptual underpinning of the *IMC Segmentation Pipeline*. We thank Nicolas Damond for providing the raw data associated to the type 1 diabetes dataset.

We also thank Daniel Schulz, Tess Brodie and Joseena Iype for critically reading and giving feedback on the manuscript.

J.W. was funded by the CRUK IMAXT Grand Challenge. B.B. was supported by a SNSF R’Equip grant, a SNSF Assistant Professorship grant, the SystemsX Transfer Project “Friends and Foes,” the SystemX grants Metastasix and PhosphoNEtX, a NIH grant (UC4 DK108132), the CRUK IMAXT Grand Challenge, and by the European Research Council (ERC) under the European Union’s Seventh Framework Program (FP/2007-2013)/ERC grant agreement no. 336921. N.E. was funded by the European Union’s Horizon 2020 research and innovation program under Marie Sklodowska-Curie Actions grant agreement No 892225.

## Notes

### Competing Interest Statement

The authors have declared no competing interest.

